# Cellular Harmonics for the Morphology-invariant Analysis of Molecular Organization at the Cell Surface

**DOI:** 10.1101/2022.08.17.504332

**Authors:** Hanieh Mazloom-Farsibaf, Qiongjing Zou, Rebecca Hsieh, Gaudenz Danuser, Meghan Driscoll

## Abstract

The spatiotemporal organization of membrane-associated molecules is central to the regulation of the vast signaling network that control cellular functions. Powerful new microscopy techniques enable the 3D visualization of the localization and activation of these molecules. However, quantitatively interpreting and comparing the spatial organization of molecules on the 3D cell surface remains challenging because cells themselves vary greatly in their morphology. Here, we introduce *u-signal3D*, a framework to assess the spatial scales of molecular organization at the cell surface in a cell-morphology invariant manner. We validated our framework by analyzing both synthetic polka dot patterns painted onto observed cell morphologies, as well as measured distributions of cytoskeletal and signaling molecules. To demonstrate the framework’s versatility, we further compared the spatial organization of cell surface signaling both within and between cell populations and powered an upstream machine-learning based analysis of signaling motifs. U*-signal3D* is open source and is available at https://github.com/DanuserLab/u-signal3D.

---

Many cell functions depend on the subcellular regulation of signal transduction. Cells regulate signaling patterns in part through nonlinear circuitry of activators and inhibitors and in part by directly controlling the localization of molecular components^2,3^. Confinement on membranes, in scaffolds, or via phase separation, all serve to localize reactions^4^. Accordingly, signaling activity varies across space and time, thus requiring live cell microscopy and appropriate image analytics for quantifying cell signaling.

One of the most important organizers of signaling states is the plasma membrane itself^5^. The plasma membrane acts as both a platform and a conduit for the chemical reactions that comprise intracellular signaling. The reactions occur at spatial scales ranging from local nano- or microscale puncta to global bursts that span the entire cell (Fig 1A)^6,7^. Deciphering the regulatory principles that define these various scales and the ensuing downstream events that control cell function requires a comprehensive quantification of molecular concentration and activity on the 3D cell membrane. This task is complicated by the fact that cells configure their plasma membranes into a wide range of convoluted 3D morphologies, for example, flattening themselves onto 2D substrates, or becoming spherical, cuboidal, or highly branched in tissues^8^. Signaling data, such as those probed by a fluorescence biosensor of protein activity, can be projected onto the 3D cell surface to facilitate analysis (Fig 1B)^1,9–11^. However, this alone does not enable quantitative comparisons of signaling organization. On the 3D cell surface, standard signal processing techniques, such as measuring spatial correlations, are stymied by the irregularly curved surface. Likewise, machine-learning approaches ranging from simple principal component analysis to complex convolutional neural networks depend on a natural ordering or parameterization of the surface signal that is consistent across morphologies, which irregularly shaped cells do not possess.

**Fig 1.**
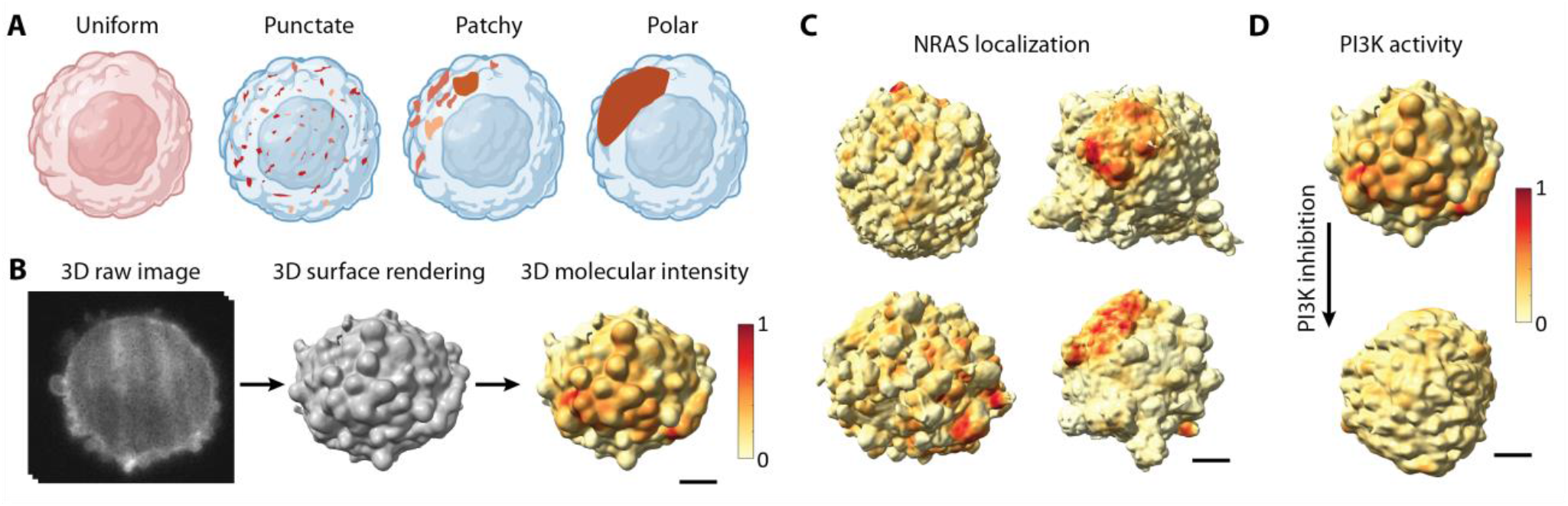
Spatial organization of molecular distributions on the 3D cell surface. **A**. Membrane-associated proteins (brown color) can spatially organize across scales ranging from a uniform distribution or punctate clustering to a highly polarized configuration. **B**. 3D raw image of an MV3 melanoma cell with PI3K activity labeled using a fluorescent biosensor (GFP-AktPH) (left), surface rendering of the same cell (middle) with molecular localization of PI3K activity projected onto the 3D cell surface (right). Red surface regions indicate high local PI3K activity. **C**. NRAS-GFP localization projected onto the 3D cell surface of MV3 melanoma cells. **D**. PI3K activity projected onto an MV3 melanoma cell before (top) and after (bottom) PI3K inhibition. All scale bars are 2 μm.

To address the challenge of ‘normalizing’ surface signal distributions across variable cell geometries, 3D analysis frameworks have considered a combination of geometry and molecular organization^12^, constrained cell shape by micro-patterned surfaces^13^, or normalized the cell shape based on global cell shape features^14–16^. Tools to normalize and analyze molecular distributions in cells adopting unconstrained morphologies are missing.

To enable the spatial analysis of the distributions of signaling and other data on 3D biological surfaces, we propose a spectral approach, in which the signaling distribution is broken down into a mathematically complete set of elementary modes that spans spatial scales. This approach relates to the spectral strategies previously used to analyze 3D cell morphologies themselves^17–19^. Whereas in these methods spectral modes were used to encode cell shape, here we exploit the modes to establish a spectral representation of the cell surface signal variation that is agnostic to the underlying cell shape variation. This allows us to i) compare the spatial organization of signaling activities across cell populations (Fig 1C), ii) measure the subcellular signaling responses to acute experimental intervention (Fig 1D), iii) perform generative modeling of representative signaling distributions, and iv) classify the features of signaling distributions across cells and conditions by machine learning pipelines.

A natural candidate for spectral decomposition of data on the cell surface is the Laplace-Beltrami operator, which constructs a mathematically complete set of Fourier-like modes on surfaces^20,21^. The Laplace-Beltrami operator has been employed to represent irregular geometries in a variety of applications, for example in biology it has been used to characterize brain morphology^22,23^. In computer science, it has been used to quantify the global characteristics of 3D shape^17,24^ and has been modified to represent local morphological motifs^25–28^. Because of its ubiquitous use in the computer graphics field, the efficiency and stability of various implementations have been extensively characterized^29–31^. Here, we use the Laplace-Beltrami operator, not to characterize shapes, but to characterize data on surfaces despite differences in the underlying surface shapes. We introduce u-signal3D, a framework to define cellular manifold harmonics and quantitatively evaluate molecular organization on the 3D cell surface across cells with highly variable morphologies. We validate our procedure by projecting synthetic signaling patterns onto experimentally measured cell surfaces of varied morphology and cell type and show the invariance of the decomposition to the underlying cell shape. To facilitate widespread use, we intuitively explain the functioning of the Laplace-Beltrami operator and provide robust implementations that fortify our workflow against the noise and artifacts typical of quantitative microscopy. Finally, to illustrate utility we apply the framework to melanoma cells, analyzing the spatial organization of cell surface signaling, cortical cytoskeletal structures, and morphological protrusions.

## Results

### Measuring the spatial organization of molecular distributions on the cell surface

Recent advances in 3D light microscopy have empowered the visualization of molecular distributions and activities at subcellular length scales^32,33^. The wealth and geometrical complexity of this data create substantial image analytical challenges. We recently contributed a software pipeline to quantify cellular morphology by extracting the cell surface from a 3D image, representing the surface as a triangle mesh, and mapping fluorescence intensities proximate to the cell surface onto the mesh itself (Fig 1B)^1^. However, since cells have different morphologies (Fig 1C), and even individual cells change their morphology over time (Supplementary Video 1) and in response to perturbations (Fig 1D) the organization of these intensity patterns cannot be directly compared across cells or time.

To orthogonalize the surface signal distributions of each cell from its morphology, we introduce a mathematical space that permits a shape-agnostic, spectral representation of the signal pattern. We employed the Laplace-Beltrami operator (LBO) to create cellular harmonics that constitute a complete set of basis functions customized to each cell morphology, implementing this approach in a software pipeline we named u-signal3D. This pipeline accepts 3D images or triangle meshes with associated local values of fluorescence intensities as the input, from which we compute spectra describing the spatial organization of surface signal distributions that can be compared across morphologies. For convenience, the pipeline includes the cell surface segmentation functionality of u-shape3D, which generates surface meshes from 3D images^1^. To meet the numerical requirements for LBO applications, we modified u-shape3D’s mesh generating process to keep only a single mesh component (Supplementary Fig.1, see Methods). The u-signal3D pipeline incorporates two implementations of the LBO, one optimized for computational speed^34^, and one robust on non-manifold meshes^30^ (Supplementary Fig. 2).

**Fig 2.**
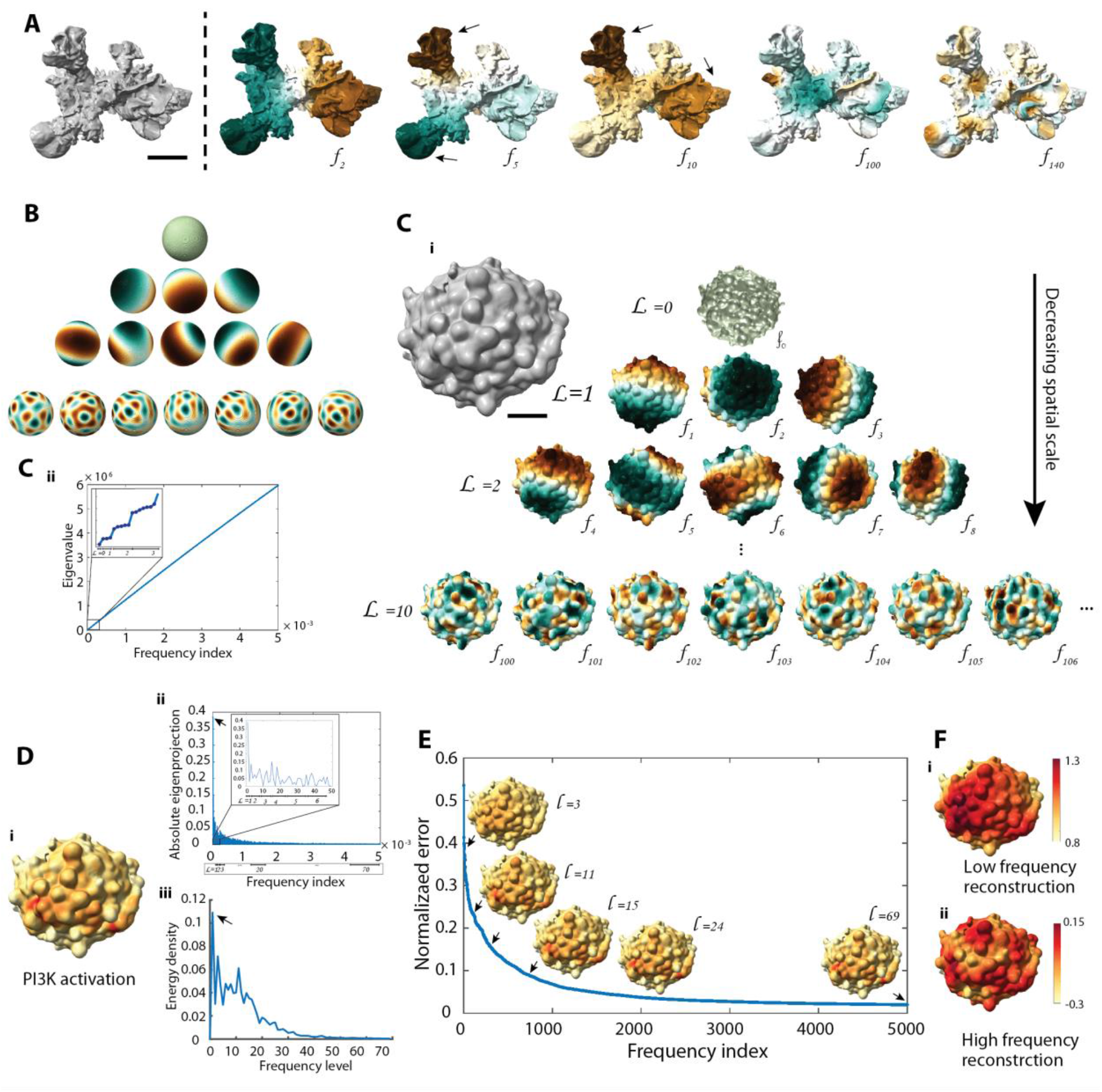
Cellular manifold harmonics for the analysis of molecular distributions on the 3D cell surface. **A**. Surface rendering of a dendritic cell (left) and selected modes of its cellular harmonics basis (*f*_*i*_) (right). Brown and green colors represent positive and negative values respectively. Black arrows indicate eigenvector magnitudes that follow protrusions in the underlying cell morphology. The cell surface was generated from the Lifeact-GFP-signal visualizing cortical actin filament structures (data not shown). Scale bar: 2 μm **B**. Some spherical harmonics, representing LBO eigenvectors applied to a sphere. The spherical harmonics naturally organize into a pyramid. **C**. Surface rendering of an MV3 melanoma cell expressing GFP-AktPH, a marker of PI3K activity. Scale bar: 2 μm. **C.i**. LBO eigenvectors (*f*_*i*_) arranged hierarchically into a pyramid from low frequency level (*L*_*j*_) to high frequency level. **C.ii**. LBO eigenvalues plotted as a function of LBO frequency index. Inset: zoomed-in region of the LBO eigenvalues plot showing only the first three frequency levels. **D.i**. PI3K activity biosensor intensity projected onto the surface of the same cell. **D.ii**. Absolute value of the LBO eigenprojection profile over the cellular harmonics index. **D.iii**. Energy density spectra of PI3K activity over frequency levels. **E**. Absolute error ratio between the reconstructed data and the original signal as a function of the number of frequency indices in the reconstruction. Black arrows indicate 3D reconstructed data visualized up to levels, L=3, 11, 15, 24, and 69. **F**. Reconstructed PI3K activity from the LBO eigenvectors showing only low frequency (L=1 to L=20) or high frequency (L=20 to L=69) information. Note color scale adjustment to visualize high-frequency variation.

The LBO computes the local geometrical variation of a 3D surface and defines a mathematically complete set of manifold harmonic functions ordered by increasing frequency (Fig 2A). These hierarchical functions conform to the morphology, for example, variations in higher order functions follow the higher geometrical variations of cell protrusions (black arrows in Fig 2A). The manifold harmonics encode simultaneously the spatial scale and orientation with respect to a coordinate frame defined by the three principal axes of cell shape. For an irregularly shaped cell, such as the branched dendritic cell shown in Fig 2A, the morphological complexity prohibits intuitive interpretations of these harmonics, especially since the orientational information dominates the scale information at higher frequencies. To illustrate the structure of the cellular harmonics, we applied the LBO to a sphere (Fig. 2B). In the limit of a perfect sphere, the LBO yields the hierarchy of the spherical harmonics, where functions at low frequency levels capture variations of the surface signals that span the entire sphere and functions at higher levels capture increasingly finer-grained variations. Each frequency level of the spherical harmonics belongs to a range of eigenvectors with similar spatial angular frequency. As an example, the basis functions at level *L* = 1 comprise three harmonics, capturing three orthogonal spatial directions. Higher level basis functions capture a greater number of pseudo-orientations with equal spatial scale. For general 3D shapes, the LBO measures the geometrical undulation of the shape about a fitting mean sphere, i.e. the sum of undulations is zero, generating a set of manifold harmonics similar to spherical harmonics. Given the similarity of the LBO eigenvectors to the spherical harmonics, we ordered the modes into a spherical harmonics style pyramid, showcasing the underlying symmetries of these cellular harmonics (Fig. 2C.i). Hence, the eigenvalues can be considered pseudo-frequencies of the geometry defined over the mesh graph and ordered inversely with the length scale of surface undulation (Fig. 2C.ii).

To quantify the organization of cell surface-associated molecular distributions, we computed the projections of these signals onto the LBO eigenvectors representing the basis functions of the underlying cell geometry. For example, Fig. 2D.i shows the fluorescence signal of a reporter of PI3-Kinase (PI3K) activity on the surface of an MV3 melanoma cell. To visualize the spatial signal content captured by the individual basis functions, we plot the function pyramid of Fig. 2C.i weighted by the magnitude of the matching eigenprojection (Supplementary Fig. 3A). This defines the contribution of each eigenvector to the total surface signal. PI3K activity has previously been shown to have a polarized signaling distribution with increased activity at the cell front and lower activity at the back^35^. Accordingly, the strongest eigenprojections associate with eigenvectors at level *L* =1 (black arrow, Fig. 2D.ii), whereas the projections for higher levels seem to decrease near-monotonically.

**Fig 3.**
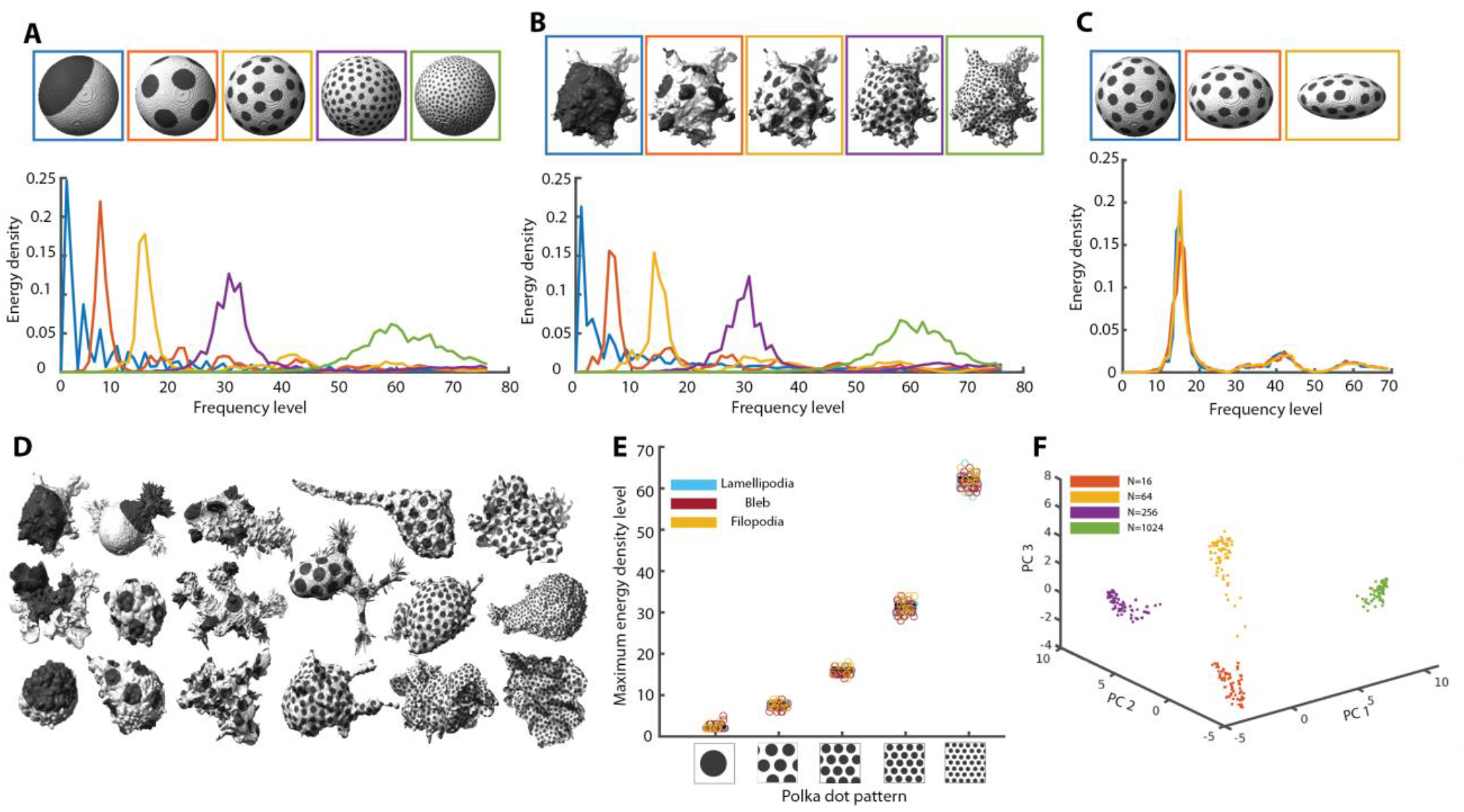
Validation and robustness of spectral decomposition analysis. **A**. Polka dot patterns on a sphere with, from left to right, dots numbering 1, 16, 64, 256, and 1024 (top). Energy density spectra for each pattern with like colors in correspondence (bottom). **B**. The same polka dot patterns as in (A) applied to the 3D cell surface of an MV3 melanoma cell expressing tractin-GFP (top) and corresponding energy density spectra (bottom). **C**. Polka dot patterns with 64 dots applied to a sphere (left) and two ellipsoids with eccentricity e = 0.75 (middle) and e = 0.5 (right) (top). Energy density spectra for each pattern with like colors in correspondence (bottom). **D**. Polka dot patterns on a variety of MV3 melanoma cells, dendritic cells and human bronchial epithelial cells (HBECs). Polka dots number 1, 16, 64, 256, or 1024. **E**. Scatter plot of the energy density peak for 67 MV3, dendritic, and HBEC cells painted with polka dot patterns of 1, 16, 64, 256, and 1024 dots. **F**. Principal component analysis of the energy

Distinct cell states are likely associated with distinct patterns of surface signaling. While this information is encoded by the mix of contributions between different spectral scales, the pattern orientation, determined by the relative weights of the basis functions at one scale, can be ignored. Seeking to separate scale and pseudo-orientational information, we turned to a quantum mechanical interpretation of the spherical harmonics. The electron orbitals of an atom are simply described by the spherical harmonics pyramid shown in Fig. 2B, with each level of the pyramid corresponding to a distinct electron energy. Indeed, mathematically all orbitals belonging to the same level have the same spatial angular frequency (Supplementary Fig. 4A). We thus separated spatial frequency from pseudo-orientation by integrating across the spherical harmonics levels. Although for non-spherical cells frequency is not strictly constant across a pyramid level, averaging out pseudo-orientational information at the rate at which it increases with frequency prevents pseudo-orientation from dominating at higher frequencies, thus allowing us to calculate an interpretable spectral signature for any given cell surface pattern.

**Fig 4.**
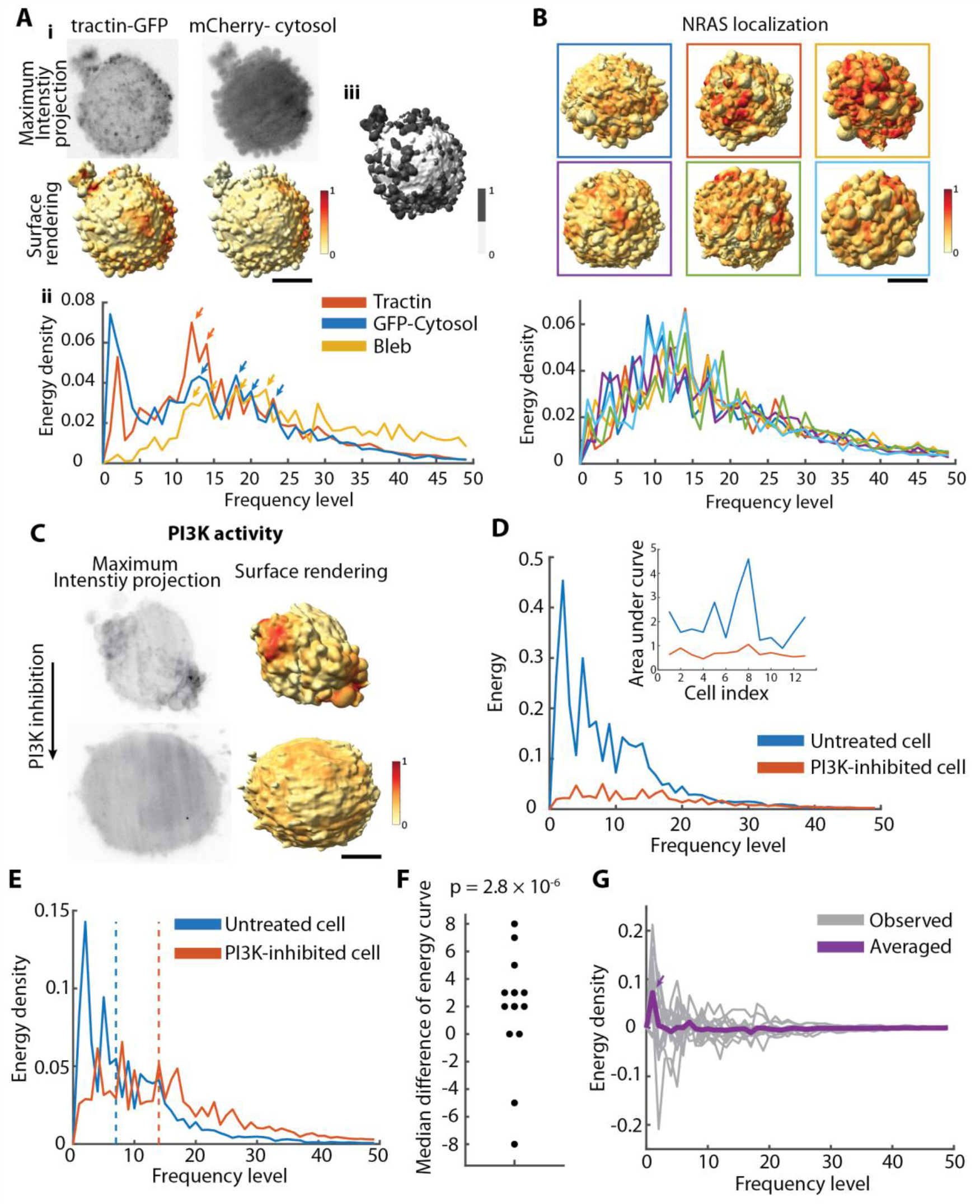
Decomposing the spatial organization of cell surface signaling on melanoma cells. **A.i**. Maximum intensity projections of an MV3 melanoma cell labeled with tractin-GFP and a cytosolic marker (mCherry) (top). Surface renderings of the same cell with fluorescence intensities projected onto the cell surface (bottom). Red indicates regions of high fluorescence intensity, whereas yellow indicates regions of low intensity. **A.ii**. Energy density spectra of tractin-GFP (red curve), a cytosolic marker (blue curve) and the binary on-/off-bleb distribution (yellow). **A.iii**. Segmentation of blebs (dark gray) vs non-bleb (white) was performed by *u-shape3D* software^1^. **B**. Surface renderings of MV3 melanoma cells expressing NRAS-GFP with fluorescence intensities projected onto the cell surfaces (top). Energy density spectra of the same cells (bottom) with like colors in correspondence. **C**. Maximum intensity projections (left) and molecular surface projections (right) of an MV3 melanoma cell before (top) and after (bottom) PI3K inhibition. The cell is labeled with GFP-AktPH, a biosensor for PI3K activity. **D**. Energy spectra of PI3K activity on the same cell before (blue) and after (red) PI3K inhibition. Inset: Area under the energy spectra curve of PI3K activity for 13 cells before (blue) and after (red) PI3K inhibition. **E**. Energy density spectra of PI3K activity before (blue) and after (red) PI3K inhibition. The dashed line indicates the mean frequency level of the energy density spectra **F**. Median energy density differences of PI3K activity between untreated and PI3K-inhibited cells (n=13 cells). **G**. The differences curve of energy spectra between untreated and PI93K-inhibited cells (gray curve is for a single cell, purple curve is averaged across 13 cells). All scale bars are 2 μm.

The overall intensity of the fluorescence signal often varies between cells for reasons unrelated to the spatial regulation of molecular concentration or activity at the surface. In particular, the total fluorescence changes as a function of the concentration of the probes used to visualize the molecular signal. To eliminate this factor, we first normalized the fluorescence intensity and further calculated an energy density across levels. This energy measures the signal gradient under Dirichlet boundary conditions, which is valid for a closed 3D shape (see Methods). The Dirichlet energy density provides a quantitative measure of how much each frequency level contributes to the organization of the surface signal. For the PI3K activity shown in Fig. 2D.i, the Dirichlet energy density reveals that the signal is organized across a range of higher spatial frequencies beyond the low level frequencies that capture front-back polarization (black arrow, Fig. 2D.iii).

Given the parameterization of a surface signaling pattern by frequency levels, we can regenerate sub-patterns containing only particular spatial scales. For example, we reconstructed the original PI3K activity pattern using LBO eigenvectors up to *L* =3, 11, 15, 24 and 69 (Fig. 2E). The absolute error ratio between the reconstructed and original signal decreases with inclusion of higher levels (Fig. 2E, Supplementary Fig. 3B), confirming that LBO eigenvectors describe the signal completely given sufficient eigenvectors. This property can be leveraged for denoising the signaling patterns by suppression of high-frequency basis functions (Supplementary Fig. 3C). Moreover, we were able to separate low frequency from high frequency components, with the former representing the signaling pattern at the cell scale, and the latter the spatial distribution of localized signaling hotspots (Fig. 2F). In summary, our LBO-based pipeline provides a cell shape agnostic means of analyzing molecular concentrations and activities at the cell surface, as well as enables filtering of surface data by spectral criteria.

### Validation of the spectral decomposition of cell surface signaling

To validate the ability of our framework to decompose cell surface signaling across spatial scales, we generated synthetic signaling patterns on both experimental and modeled cell geometries. The frequency-dependent behavior of optical instruments, including microscopes and telescopes, is commonly assessed via stripe patterns, such as Ronchi rulings^36^. Stripe patterns cannot be applied to the highly curved surfaces of cells with any consistency. We instead introduced polka dots consisting of circular shapes of equal radius arranged as far apart as possible on the cell surface (Supplementary Fig. 5, see Methods). Generating variously scaled polka dot patterns allowed us to simulate surface signaling distributions ranging from front-back polarization to punctate micro-clusters (Fig. 3A). Applied to a sphere, coarse-to fine-grained polka dot patterns yielded distinct spectral peaks at increasingly higher frequency levels (Fig. 3A). The peaks broaden at higher frequencies, in part because although the spherical harmonics have the same spatial angular frequency at each pyramid level, the number of peaks varies (Supplementary Fig. 6A, Supplementary Video 2). At higher frequency levels, similar peak numbers are shared by a greater range of levels (Supplementary Fig. 6B, Supplementary Video 3). Consequently, polka dot number and angular spatial frequency are not perfect analogues, even though both capture spatial scale information.

Analyzing the same polka dot patterns on a melanoma cell surface, we recovered very similar spectra (Fig. 3B), confirming that the spectral decomposition is largely independent of the underlying cell morphology. To more formally test the consistency of spectra across cell morphologies, we first applied polka dot patterns to a sphere and similarly sized ellipsoids (Fig. 3C). The spectra of any given polka dot pattern peaked at the same frequency level. We then tested the robustness of the spectral decomposition across a cell library encompassing different cell types with different morphological motifs (Fig. 3D, Supplementary Fig. 7)^1^. The motifs include lamellipodia (thin, sheet-like membrane protrusions), blebs (hemispherical membrane protrusions), and filopodia (elongated and spiky membrane protrusions). For five polka dot patterns, we found consistent spectral peaks over 67 different cell morphologies (Fig. 3E). Our spectral decomposition thus characterizes the spatial scales of cell surface patterns in a cell shape-agnostic manner.

Measuring cellular harmonics not only allows us to characterize the spatial organization of cell surface signals, but also to parameterize them for downstream analyses such as machine learning applications. As an example, we used principal component analysis to reduce the dimensionality of the energy density spectra of the abovementioned five polka dot patterns on 67 cell morphologies (Fig. 3F). Independent of the cell morphology, each polka dot scale defines a distinct cluster in the reduced space. Of note, performing a similar cluster analysis directly on the intensities projected onto the cell surface is impossible, because there is no natural ordering of intensities that is consistent between cells with highly diverse morphologies.

In summary, our polka dot validation established the ability of the pipeline to measure the spatial organization of cell surface patterns independent of the underlying cell shape.

### Spectral decomposition of cytoskeletal organization and signaling events on melanoma cell surfaces

To illustrate the biological utility of cellular harmonics, we applied our framework to the analysis of molecular distributions in melanoma cells imaged in 3D culture via high-resolution light-sheet microscopy^35,37,38^. As a first example, we compared the cell surface distributions of two co-imaged molecular signals. Melanoma cells in soft collagen tend to extend dynamic blebs. Expanding blebs are thought to be largely devoid of cortical actin whereas retracting blebs have increased cortical actin^39^. Thus, a snapshot of the cortical filamentous actin distribution at a single time-point of a blebbing cell is expected to display a patchy fluorescent signal over the entire cell surface (Fig. 4A.i and Supplementary Video 4). We measured the localization of filamentous actin (tractin-GFP) and a cytosolic GFP marker within a 1 μm sampling radius of the cell surface. Aligned with our expectation, filamentous actin shows peaks in spatial organization at mid-range energy levels (red arrows in Fig. 4A.ii), whereas the cytosolic marker has more uniform spatial organization (blue arrows in Fig. 4A.ii). Of note, the first peak in both cytosol and tractin energy density spectra is associated with an illumination gradient across the entire cell, which is clearly recognizable in the maximum projection images (Fig. 4A.i).

To enable interpretation of energy spectra, we calculated an approximate conversion between energy level and actual spatial scale. To do so, we measured the relative distances amongst polka dots distributed on these blebby cells (Supplementary Fig. 8). Comparison between the spectra of tractin-GFP and polka dot patterns suggests that filamentous actin is organized at a peak length scale of 6 to 8 μm (Supplementary Fig. 8, blue box). Moreover, these analyses indicated that 50 frequency levels are sufficient to represent energy density spectra at a spatial resolution above 1 μm, which is the limit imposed by our choice of a sampling radius to compute cell surface signals (Supplementary Fig. 8, see Methods). To test the possibility that the spatial pattern of actin filaments relates to the spatial pattern of blebs, we detected blebs using the u-shape3D software and converted the segmentation into a binary cell surface pattern (Fig. 4A.iii). The energy spectra of bleb distribution peaks (L= 10 to 25) at a spatial scale of 4 to 8 μm (Supplementary Fig. 8, purple box). Collectively, these measurements indicate that bleb and filamentous actin density are organized, as expected, at similar spatial scales.

Next, we analyzed the spatial organization of wild type NRAS signaling in melanoma cells. Across six cells, the NRAS organization peaked at mid-range energy levels (L=10 to 15) corresponding to spatial scales of 7 to 9 μm (Supplementary Fig. 9, blue box). NRAS is recruited to the membrane by bleb-related scaffold proteins^37^. Thus, we hypothesized that the spatial scale of NRAS signaling relates to the length scale of bleb-induced surface undulations. Of note, although we established that the energy spectra of the spatial distributions of cell surface signals are shape-agnostic, the signal distributions themselves may very well be coupled to the spatial distribution of morphological motifs. We detected blebs on the cell surface using the u-shape3D software (Supplementary Fig. 10). Compared to the bleb spectra, the NRAS spectra were somewhat skewed to lower frequency levels, which captures whole-cell scale polarity. Thus, we concluded that while the bleb distribution in these melanoma lacks an axis of preferential organization, the NRAS distribution is determined by a secondary, long-range mechanism inducing cell-scale polarity.

Polarized cell signaling is often accompanied by polarized cell morphology. Our framework allows the disentangling of morphology-driven and signaling-driven polarization. For example, PI3-Kinase activity is known to be polarized in migrating melanoma cells that themselves have polarized morphologies^35^. We measured PI3K activity via a fluorescent biosensor (GFP-AktPH) both before and after treatment with a PI3K inhibitor (PI3K*α* Inhibitor). Even when the effects of a polarized morphology were excluded, PI3K activity was polarized as apparent from the high energy at low frequency levels. Upon PI3K inhibition, the overall energy spectra is reduced (Fig. 4D), indicating that the spatial organization of the PI3K signal is governed at least in part by the PI3K activation level, i.e. via chemical feedbacks. The area under the energy curve for 13 cells demonstrates the energy reduction after PI3K inhibition for individual cells (Fig. 4D inset). Moreover, even on a relative scale the energy density at low frequencies is suppressed, showing that PI3K inhibition abrogates polarity in signaling (Fig. 4E). Calculating the mean frequency level of energy density spectra (dashed lines in Fig. 4E) before and after treatment for 13 cells and applying the Kolmogorov-Smirnov test on the difference between the mean frequency level before and after the treatment, we found that PI3K inhibition significantly raised the average frequency level of the energy density distribution, indicating that the signaling was less polarized (Fig. 4F), consistent with the qualitative conclusions drawn in^35^. We also computed the averaged difference between the curves before and after treatment over 13 cells (Fig 4G). The biggest difference occurs at low frequency levels, confirming that PI3K inhibition abrogates primarily the signaling polarity (purple arrow in Fig 4G).

## Discussion

Elucidating the organization of molecular distributions and signaling activities on cell surfaces in 3D is confounded by the complexity of relating surface patterns to a reference that allows comparative analyses across cell populations with widely variable morphologies. Here we introduce cellular harmonics, a method that disentangles the organization and heterogeneity of cell surface signaling from the heterogeneity of cell morphology. We implemented the Laplace-Beltrami operator (LBO) on individual cell shapes to define a complete basis that spans spatial frequencies and is well adapted to the morphological structures of the cell surface. To parameterize a molecular cell surface pattern, we generate spectra that capture the spatial scale signatures of the pattern. Using polka dot patterns, we validated that the spectra reflect the scales of surface signal organization independent of the underlying cell shape. We employed the proposed framework to analyze fluorescence microscopy images of molecular distributions on the cell surface, in particular, comparing molecular organization across a cell population, and evaluating the molecular redistribution in the presence of an acute drug treatment. Finally, we illustrated that the generated spectra define an informative feature set for further processing in machine learning-based classification of molecular patterns.

Unlike previous approaches to quantify molecular cell surface patterns in the face of cell morphological variation, which relied on a few global features to represent cell shape or constrained cell shape on micro-patterns ^13–15^, our work introduces a mathematical construction to characterize the signal organization in a functional space that captures the full range of natural scales relevant to the regulation of molecular processes. Thus, our framework is suitable to experimental conditions, including *in vivo* imaging, that preclude external control over cell shape. In general, the spectral decomposition of surface data enables the measurement and interpretation of the spatial organization of molecular distributions ranging from simulated data, such as signaling distributions on the cell surface, to larger scale phenomena, such as cytoskeletal distributions on embryo surfaces. The framework also provides a mathematically tractable, yet complete, representation to power downstream analyses including generative modeling and machine learning.

The u-signal3D framework is made available as a set of MATLAB functions bundled into a user-friendly interface. Underlying the implementation of u-signal3D is a robust computational pipeline for mesh triangulation and determination of LBO eigenvectors using algorithms that cope with both manifold and non-manifold cell geometries ^30,34^. The Laplace-Beltrami operator is critical to many computer graphics algorithms. The resources and validation provided by this work will aid the microscopy community in further adapting tools from the computer graphics field.

## Methods

### Cell culture and imaging

To validate our workflow, we used movie collections that were generated for previously published studies of various cellular processes. All cells were imaged via high resolution light-sheet microscopy^38,40^. The cell preparation and imaging conditions for the tractin-GFP^38^, NRAS-GFP^37^, and PI3K activity^35^ (GFP-AktPH) datasets are available in the previously published papers. To simulate the polka dot patterns on observed cell geometries, we also used 67 cell surfaces from Driscoll *et al*^1^. These cell surfaces included 29 dendritic cells expressing Lifeact-GFP, 27 MV3 melanoma cells expressing tractin-GFP, and 11 human bronchial epithelial (HBEC) cells expressing tractin-GFP.

### Cell surface generation

The pipeline embeds the cell surface generation modules from our previously published u-shape3D analysis framework^1^. First, 3D raw images of cells were deconvolved using either a Wiener filter with the Wiener parameter set to 0.018 (for tractin-GFP and NRAS-GFP localization) or a Richardson-Lucy algorithm (for PI3K activity). To reduce deconvolution artifacts, images were apodized, as previously described^38^. Then we smoothed the deconvolved images with a 3D Gaussian kernel (only for the PI3K dataset with *σ* = 1.5 pixels) and applied a gamma correction (for the PI3K dataset of 0.7, for the NRAS and tractin-GFP datasets of 0.6). For the segmentation of melanoma cells in the PI3K and NRAS data sets, we used the ‘twoLevelSurface’ segmentation mode, which combines a blurred image of the cell interior with an automatically thresholded image of the cell surface. For melanoma cells labeled with tractin-GFP, cells were segmented at the intensity value specified by the Otsu threshold^41^. Finally, the triangle mesh representing the cell surface was smoothed using curvature flow smoothing^42^.

3D surface renderings of single cells were created with ChimeraX^43^ and schematics were created with BioRender.com.

### Mesh characterization and validation

Triangle surface meshes generated from 3D microscopy images are prone to artifacts that adversely impact the Laplace-Beltrami operator output. One such artifact is a multi-component mesh, which often has isolated sets of disconnected triangles that do not correspond to the actual cell geometry (Supplementary Fig. 1, black arrows), and thus can be safely removed. Connected components of a mesh are identified by a list of mesh faces that share an edge. Smaller components are filtered out by a threshold defined on volume or surface area. For our data, we set the threshold to 10% of the total volume, which is sufficient to generate a cell surface with a single component. We used the *‘remove_small_components’* function from a public GitHub repository of the geometry processing toolbox [https://github.com/alecjacobson/gptoolbox].

For further processing, the cell surface must be represented as a closed 2-manifold mesh satisfying two conditions: i) each edge is incident to only two faces and ii) at each vertex of a face the face is embedded in a fan (Supplementary Fig. 2B). Before computing the Laplace-Beltrami operator, we checked whether the mesh surface was manifold or non-manifold. If it was manifold, we used a standard approach to compute the LBO (next section). If there were non-manifold vertices, we used a state-of-the-art LBO algorithm^30^ that first creates a manifold mesh, termed a tufted mesh, from the non-manifold mesh, before applying the LBO.

### Intensity measurement

We measured the fluorescence intensity local to each mesh vertex from the raw image. The cytosolic background noise was first removed by subtracting the median pixel intensity inside the cell. Then, at each vertex, the average pixel intensity was computed within a sampling radius of 1 μm over the pixels inside the cell. Finally, we normalized each cell’s surface intensity to a mean of one.

### Laplace-Beltrami calculation

The cell surface geometry is defined by vectors ***X*** = (*x, y, z)* of *N* triangle mesh vertices in a Cartesian coordinate system. The mesh vertices are connected in triangles to form mesh faces. To create a set of basis functions for a cell, we applied the Laplace-Beltrami operator (LBO) to the mesh. The LBO is a second-order differential operator defined as the divergence of the gradient of a function. On a triangle mesh, it computes the geometrical variation of the surface from a sphere. Mathematically, we aim to solve the following eigendecomposition equation to obtain the eigenvectors and eigenvalues of the LBO on a given mesh:

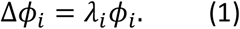

Here *ϕ*_*i*_ denotes the *i*-th eigenvector corresponding to the *i*-th eigenvalue, *λ*_*i*_, of the Laplacian operator, Δ.

The Laplacian operator acting on an arbitrary discrete mesh has been extensively studied^20,29,34,42,44–46^. Most computational approaches are based on the cotangent scheme^34^, which is a form of first order finite element method applied to the Laplacian operator on a surface. In the cotangent formula (*C*), the LBO is an *N*x*N* symmetric matrix:

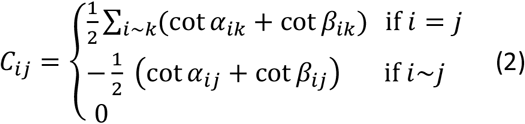

where *α*_*ij*_, *β*_*ij*_ are the two angles relative to edge *ij* as indicated in Supplementary Fig. 2A. For the matrix element *C*_*ii*_, the index *k* runs over all vertices connected to vertex *i* through an edge *ik*.

To obtain the LBO eigenvectors and eigenvalues, we used the function *eigs*() in MATLAB. The eigenvectors are frequency-based hierarchical functions that follow the protrusions of the 3D shape, respecting their symmetries. For example, for a sphere the eigenvalues of the first frequency level (*L* =1) are identical and correspond to three eigenvectors oriented in three orthogonal directions in 3D space (Supplementary Fig. 4A). When the LBO is applied to an ellipsoid, where the symmetry is broken in one direction, the eigenvalue has higher magnitude in the short ellipsoid axis direction (Supplementary Fig. 4B). Eigenvectors at the higher frequency levels follow the symmetry breaks of the ellipsoid to capture the shape variation relative to a sphere. In analogy, for a complex shape, the LBO eigenvectors follow the variation of the surface relative to a sphere (Fig 2C).

### Representation of cell surface signals using LBO eigenfunctions

Eigenvectors (*ϕ*_*i*_) of the Laplacian constitute an orthogonal complete basis set. Thus, we can extend a surface signal *u*(***X***) in this basis,

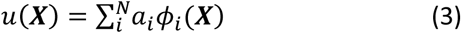

where *N* is the number of Laplacian eigenvectors and *a*_*i*_ is the Laplacian eigenprojection of the *i*-th eigenvector. The eigenprojections determine the contribution of each cell geometry defined eigenvector in parameterizing the signal *u*(***X***). Based on the LBO eigenvectors and eigenprojections of a signal, the signal can be recreated 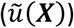. To do so, we summed up the eigenprojections, multiplying each by the corresponding eigenvector in order to reconstruct the signal. To assess the reconstructed signal with its original values, we computed the absolute error ratio given by,

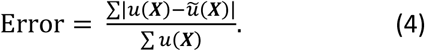

### LB frequency level measurement

Application of the LBO (equation (1)) to a perfect sphere yields the spherical harmonics, *Y*_*l,m*_(*θ, φ)*, as shown in a frequency level pyramid with the first levels shown in Fig. 2B. In spherical coordinates, the spherical harmonics are given by

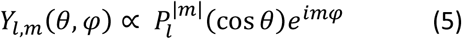

in which, *θ, φ* define the colatitude and longitude, respectively; the indices *l* and *m* indicate the degree and order of the function in frequency-based levels; and 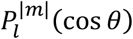 are the Legendre polynomials. The LBO encodes both the spatial angular frequency and its orientation over the eigenvectors. Each frequency level has 2*l*+1 eigenvectors. As an example, the second level (*l* =1) of spherical harmonics has three eigenvectors with identical spatial angular frequency (Supplementary Fig. 4A). To quantify the spatial angular scale of a signal on a mesh surface, we collapsed the measurements from individual eigenvectors (frequency index) to a single value per level (frequency level), as we explain in the next section. Although for complicated 3D shapes with lower symmetry compared to a sphere, eigenvectors at the same frequency levels have different eigenvalues, it is still reasonable to smooth the measurement over each level (*l*) to extract the spatial scale signature of a given molecular organization.

### Dirichlet energy calculation

In computer geometry and shape analysis, the Dirichlet energy is often used as a metric of the smoothness of a function defined in a region^47^. The Dirichlet energy is given by

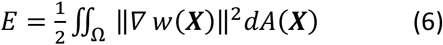

where *w*(***X***) is a function defined on the region *Ω* in three dimensions ***X*** = (*x, y, z*). Integration by parts yields,

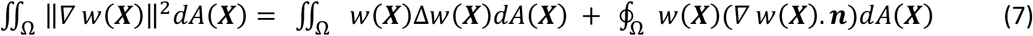

The first term on the right hand side is the Laplacian of the *w*(***X***) in the region, and the second term relates to the boundary conditions. For a closed surface, such as a segmented cell, this term vanishes.

Substituting *u(****X***) for *w(****X***) subject to equation (1) and under consideration of the orthogonality of the LBO eigenvectors, the energy term equation (7) turns into,

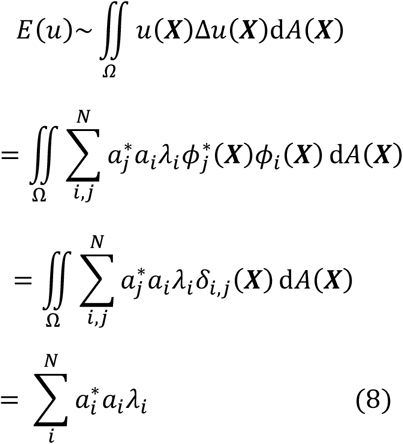

The Dirichlet energy in (8) is a scalar parameter. In our framework, the energy is computed per spatial angular frequency level to obtain an energy spectra. The energy per frequency level follows

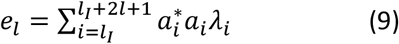

where *l*_*I*_ is the first index of the eigenvector at frequency level *l*. Similar to quantum physics, the 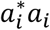 term gives the probability of the *i*-th eigenvector contributing to the signal function *u*(***X***), while *λ*_*i*_ is the eigenvalue corresponding to the eigenvector. As mentioned above, the eigenprojections per level indicate the dominant spatial scale of a given molecular organization without regard to shape symmetry variation. Since fluorescence expression can vary from cell to cell, we normalized the energy spectra to 1 by dividing by the total energy (*E* in equation (8)) to define an energy density spectra for each cell that can be compared across a cell population.

### Polka dot generation

We simulated polka dot patterns to validate the spatial scale of the measured signal distribution on a given mesh. As shown in Supplementary Fig. 5, to generate the polka dots, dot centers, or seeds, were first found by maximizing the minimum pairwise Euclidean distance amongst seeds. We used the ‘*farthest_point’* function from a public GitHub repository of the geometry processing toolbox [https://github.com/alecjacobson/gptoolbox]. Then we calculated the geodesic distance of all vertices from the closest seed using the fast marching method. The polka dot pattern was then created by thresholding the surface over the list of geodesic distances, such that the fraction of total surface area occupied by polka dots was maintained across patterns. For presented data, we set the polka dot surface area to 30% of the entire cell surface area.

### Polka dot clustering

We constructed four polka dot patterns with 16, 64, 256, and 1024 dots respectively on each of the 67 cell morphologies. For each pattern, we compiled a feature vector composed of the energy density spectra and then performed principal component (PC) analysis. The first three components contributed 58% of the total variance in the 77-dimensional feature space. To cluster the polka dot patterns, we employed the k-means algorithm in the 3D PC space. The number of clusters was set to four, the number of polka dot patterns.

### Statistical analysis of PI3K inhibition

To quantify the redistribution of PI3K activity after PI3K inhibition, we compared the energy spectra of individual cells before and after inhibition. We measured the mean frequency level of each curve and computed the difference in the mean before and after inhibition for 13 cells (Fig. 4E,F). Additionally, to evaluate the energy spectra over the entire frequency range, we averaged the difference curves between the before and after inhibition conditions for the 13 cells (Fig. 4G).

## Data Availability

The data that support the findings of this study are available from the corresponding author upon reasonable request.

## Code Availability

All analysis was performed in a custom software package (u-signal3D) written in MATLAB (Mathworks), which is available at [https://github.com/DanuserLab/u-signal3D].

## Acknowledgments

We would like to thank Maks Ovsjanikov, Jungsik Noh, and Jaewon Huh for fruitful discussions. This work was supported by the following grants: K99GM123221 to MKD and R35GM136428 to GD.

## Supplementary figures and movies

**Fig S1.**
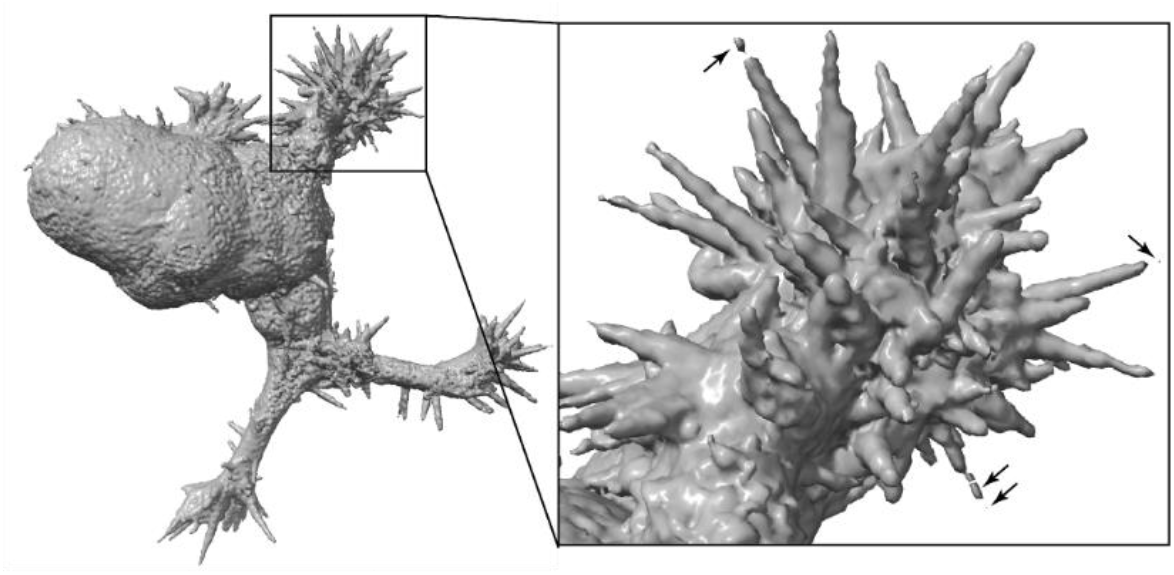
Multi-component triangle mesh. Surface renderings of a human bronchial epithelial cell (HBEC). The cell surface was generated from the Lifeact-FP-signal visualizing cortical actin filament structures (data not shown). Multiple mesh components are visible. They are automatically filtered by the *u-signal3D* software.

**Fig S2.**
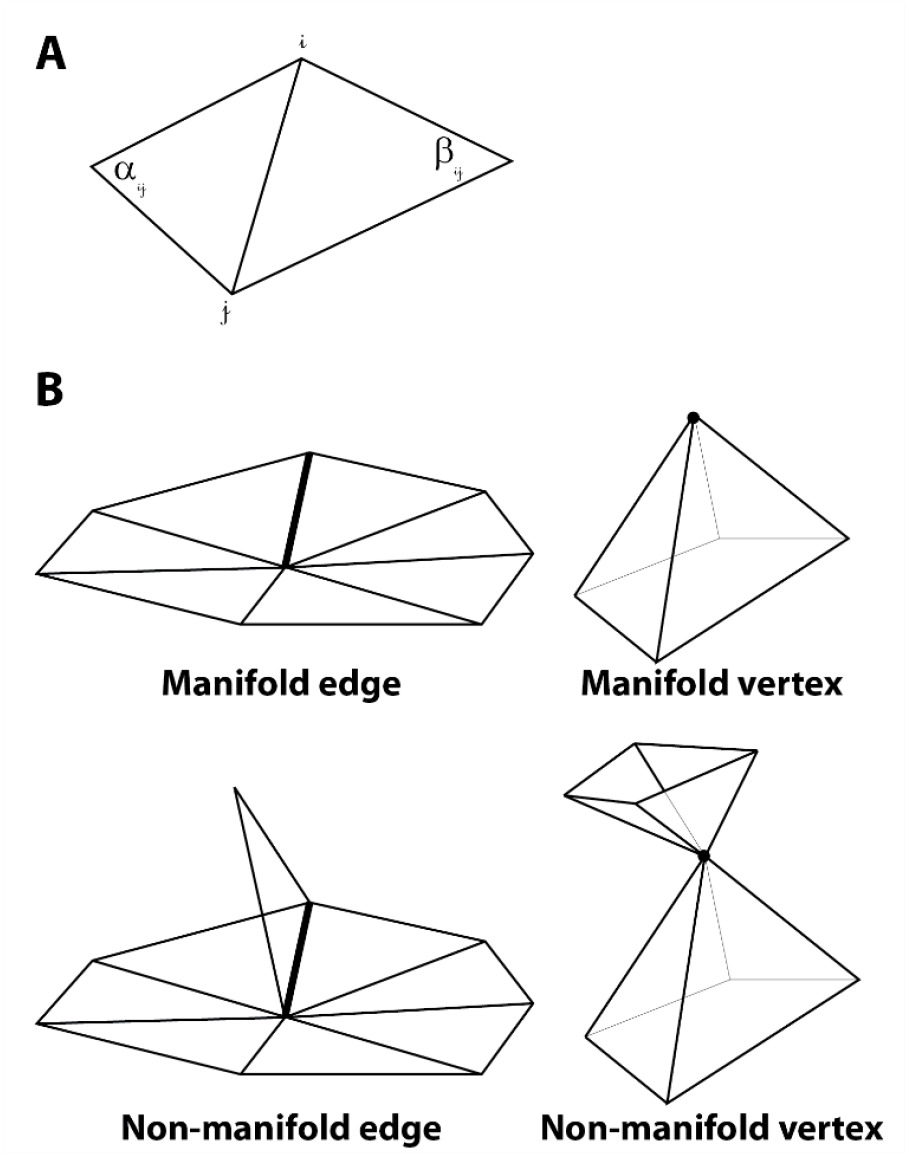
Manifold versus non-manifold meshes. **A**. Two faces of a triangle mesh. α_ij_and β_ij_ are the angles corresponding to the *i*-th and *j*-th vertices in the cotangent Laplacian method. **B**. Manifold and non-manifold edges and vertices in a triangle mesh.

**Fig S3.**
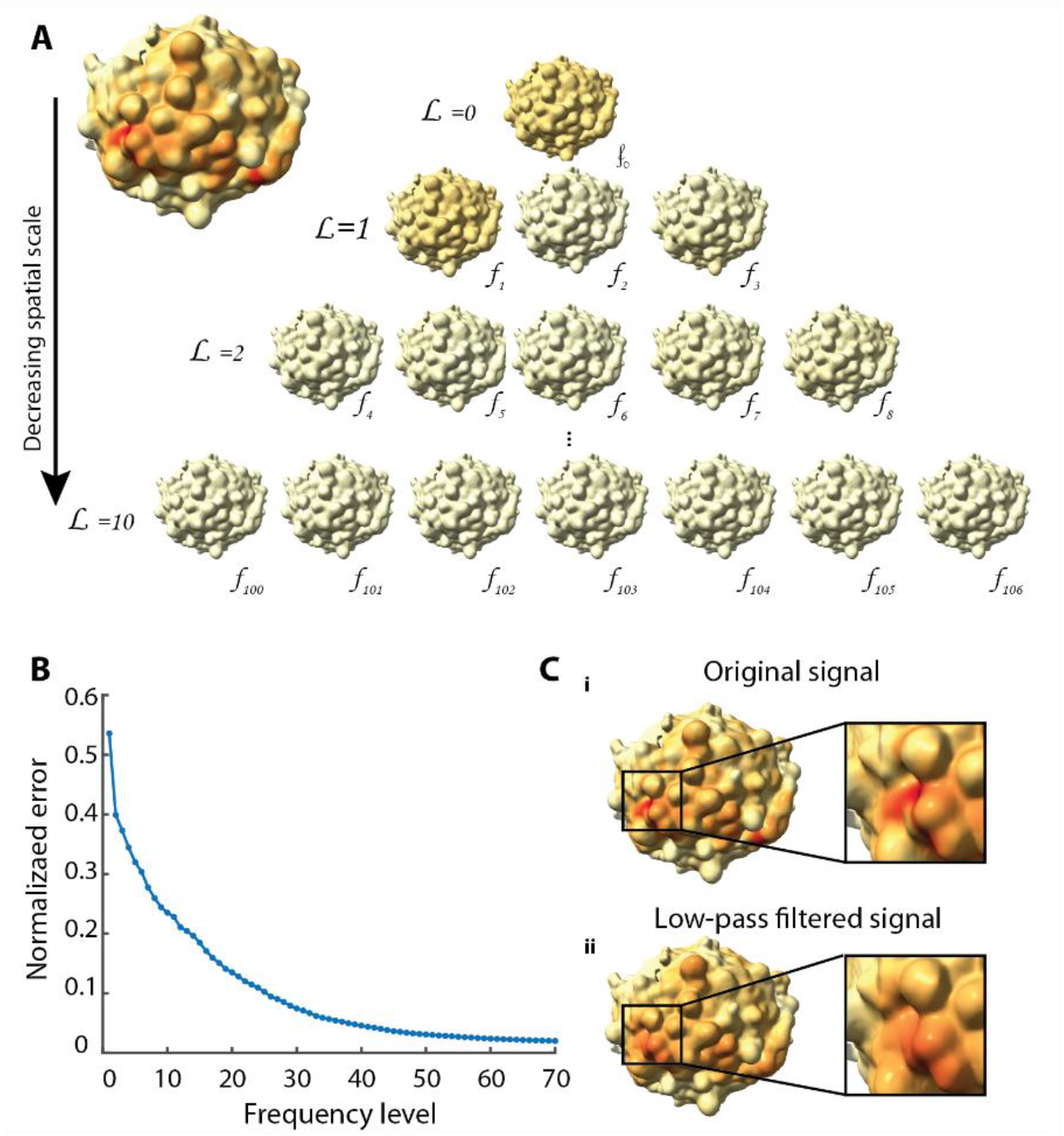
Reconstructing surface signal data from the spectral decomposition. **A**. PI3K activity biosensor intensity (GFP-AktPH) projected onto MV3 melanoma cells. LBO eigenvectors (*f*_*i*_) corresponding to the shown PI3K activity are organized hierarchically from low frequency level (*L*_*j*_) to high frequency level. **B**. Absolute error ratio between the reconstructed surface signal distributions and the original PI3K activity biosensor intensity as a function of the number of frequency levels included in the reconstruction. **C**. PI3K activity (GFP-AktPH) projected onto the same MV3 melanoma cell with the original data (**i**) shown compared to the low-pass filtered signal up to level L=10 (**ii**).

**Fig S4.**
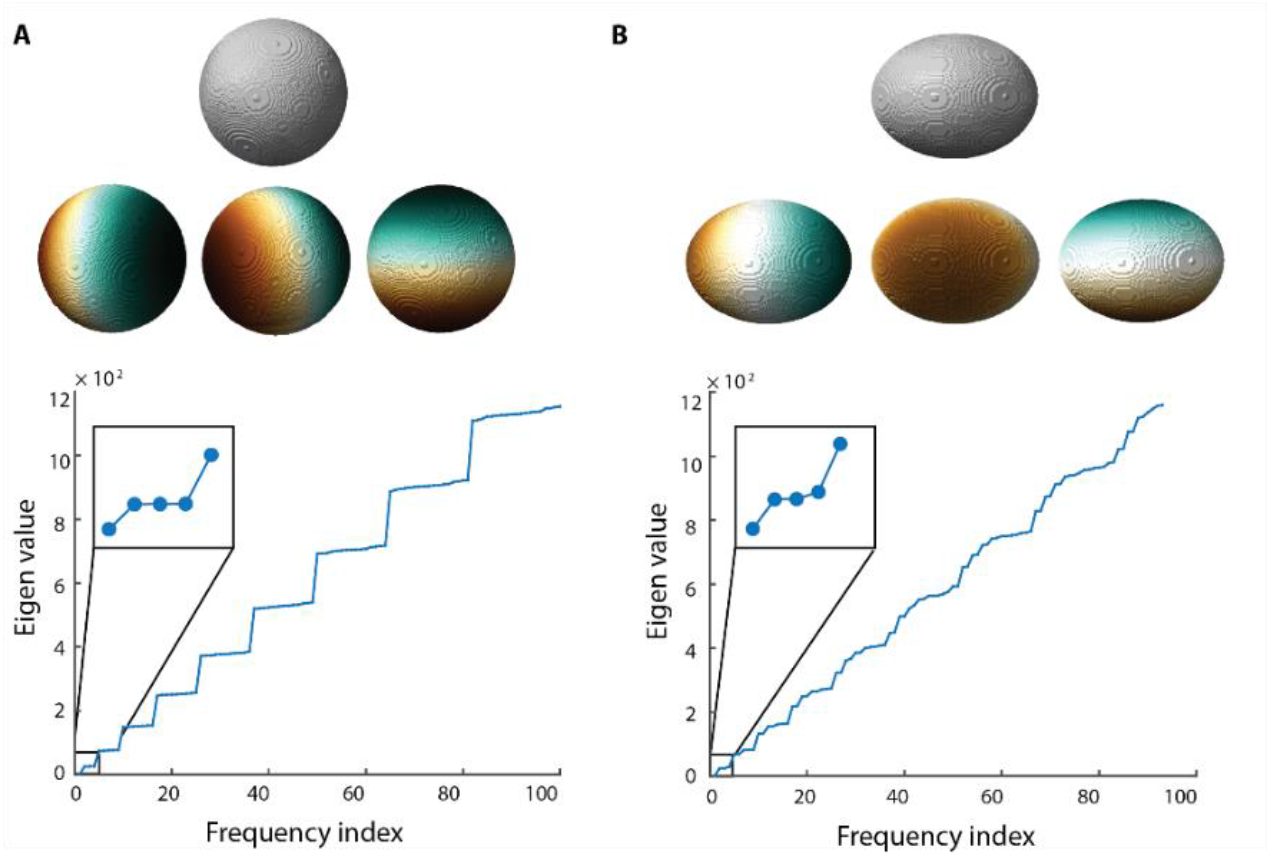
The Laplace-Beltrami operator (LBO) on symmetric shapes. **A**. LBO eigenvectors of level L=1 applied to a sphere (top). Brown and green colors represent positive and negative values, respectively. LBO eigenvalues of the sphere (bottom). Inset: zoomed-in region of the eigenvalue graph showing low frequency levels. **B**. LBO eigenvectors of L=1 applied to an ellipsoid with e=0.75 (top). Brown and green colors represent positive and negative values, respectively. LBO eigenvalues of the ellipsoid (bottom). Inset: zoomed-in region of the eigenvalue graph showing low frequency levels.

**Fig S5.**
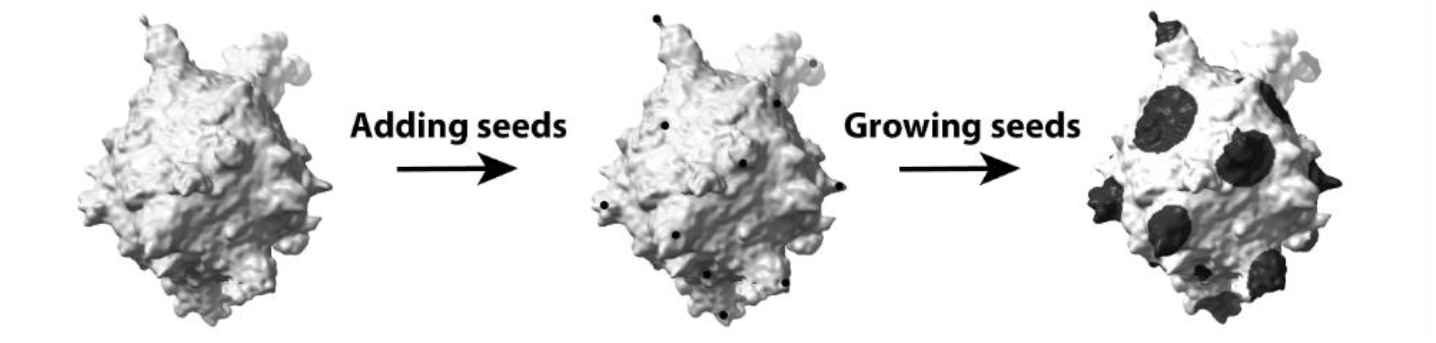
Polka dot pattern generation. Surface rendering of an MV3 melanoma cell expressing tractin-GFP (left). The same cell surface painted with 64 dots (middle), and grown to 64 polka dots with surface area totaling 30% of the entire cell surface area (right).

**Fig S6.**
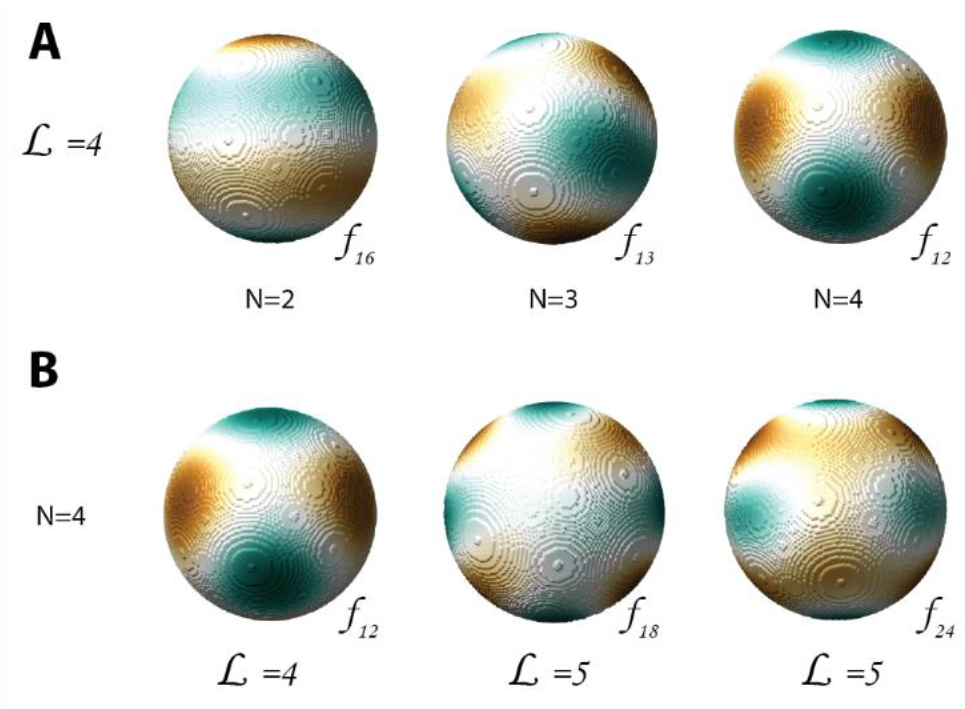
Number of LBO eigenvector peaks vary even at the same frequency level. **A**. Examples of LBO eigenvectors of a sphere at frequency level L=4. The number of peaks in the eigenvector magnitude is 2 (left), 3 (middle), and 4 (right). **B**. Examples of LBO eigenvectors of a sphere that have 4 peaks in magnitude, at frequency levels L = 4 (left), 5 (middle), and 6 (right).

**Fig S7.**
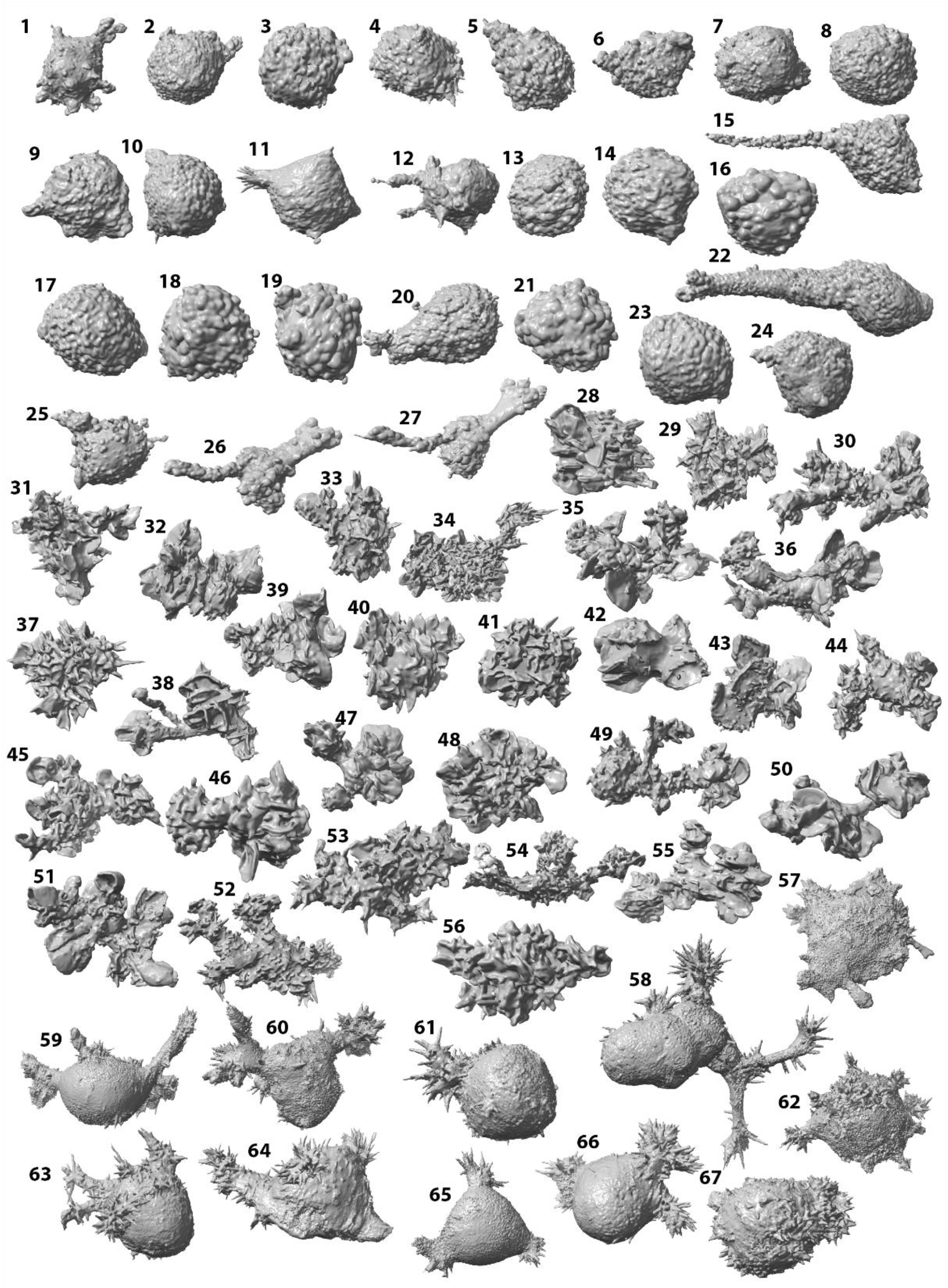
Cell surface library. Surface renderings of MV3 melanoma cells (1-27), dendritic cells (28-56), and HBEC cells (57-67). The cell surfaces were generated from the tractin-GFP intensity (MV3 cells, HBEC) and Lifeact-GFP (dendritic cells).

**Fig S8.**
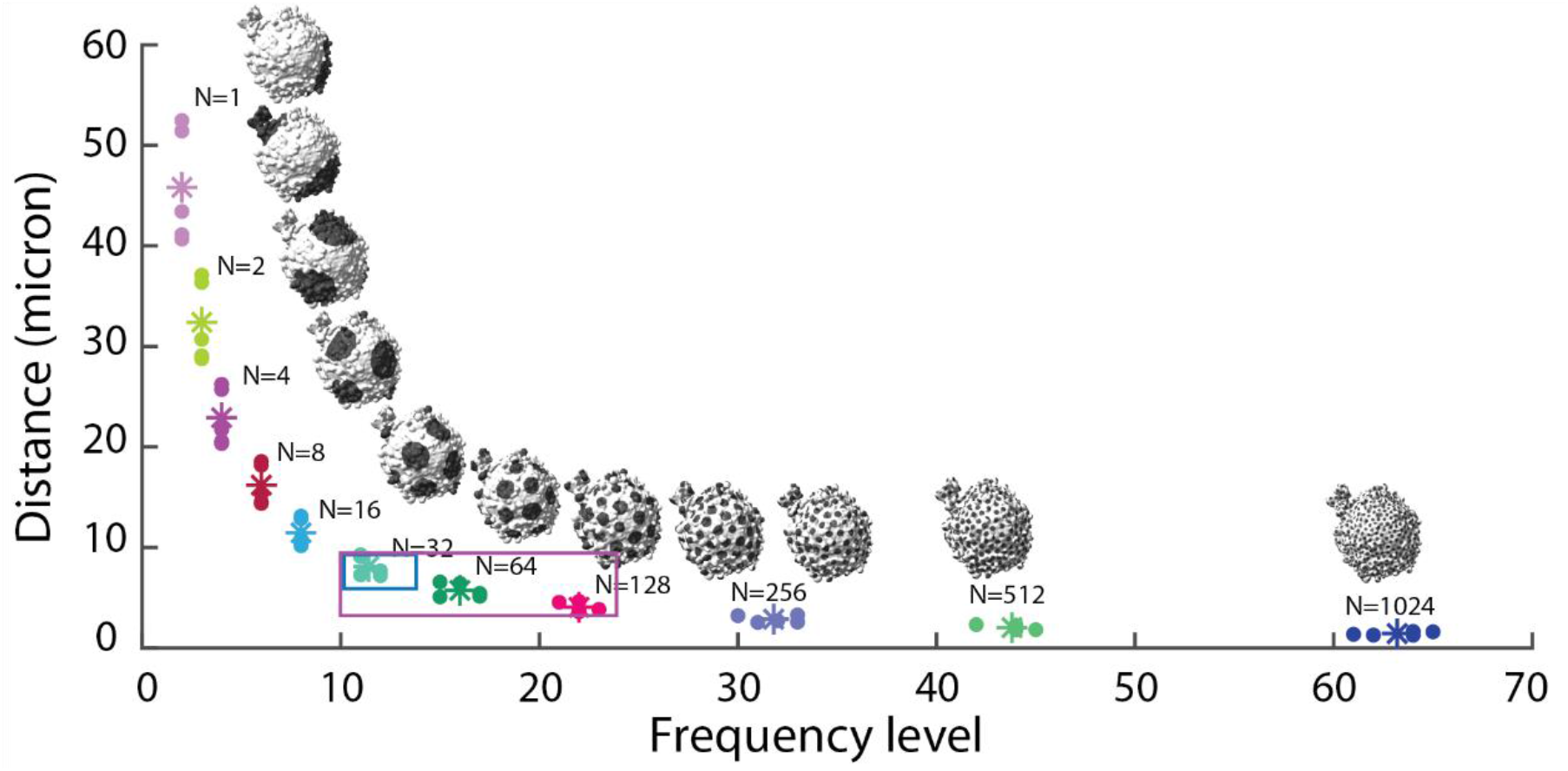
Distance conversion of LBO frequency levels on tractin-GFP labeled melanoma cells. Geodesic distance between the polka dots on a mesh surface (presented in Fig. 4A) as a function of energy peak level. The polka dots painted on each mesh number 1, 2, 4, 8, 16, 32, 64, 128, 256, 512, or 1024. The cell surface were generated from the cytosolic marker (mCherry) intensity.

**Fig S9.**
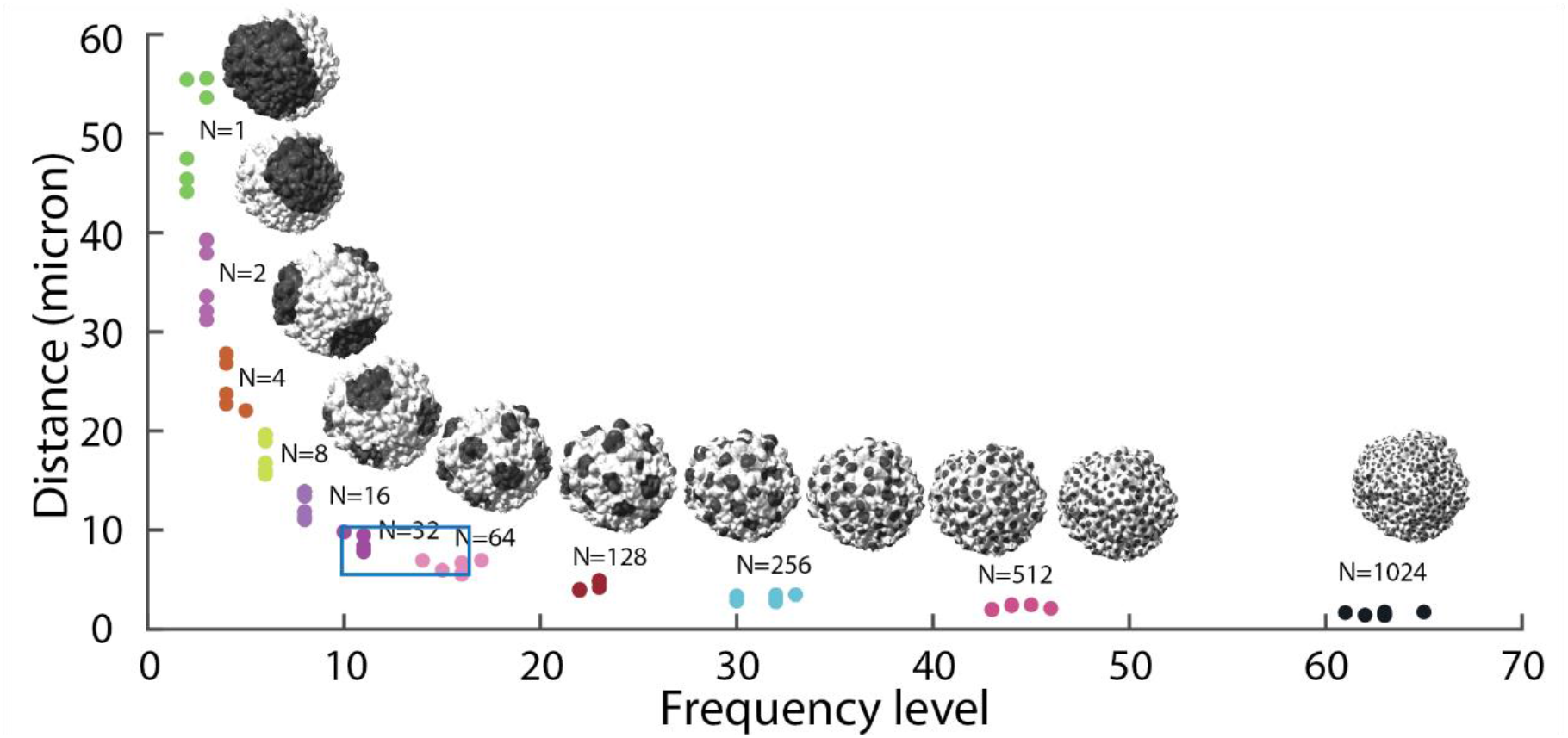
Distance conversion of LBO frequency levels on NRAS-GFP labeled melanoma cells. Geodesic distance between the polka dots on a mesh surface as a function of energy peak level for an MV3 melanoma cell (presented in Fig. 4B). The polka dots painted on each mesh number 1, 2, 4, 8, 16, 32, 64, 128, 256, 512, or 1024. The cell surface were generated from the NRAS-GFP intensity.

**Fig S10.**
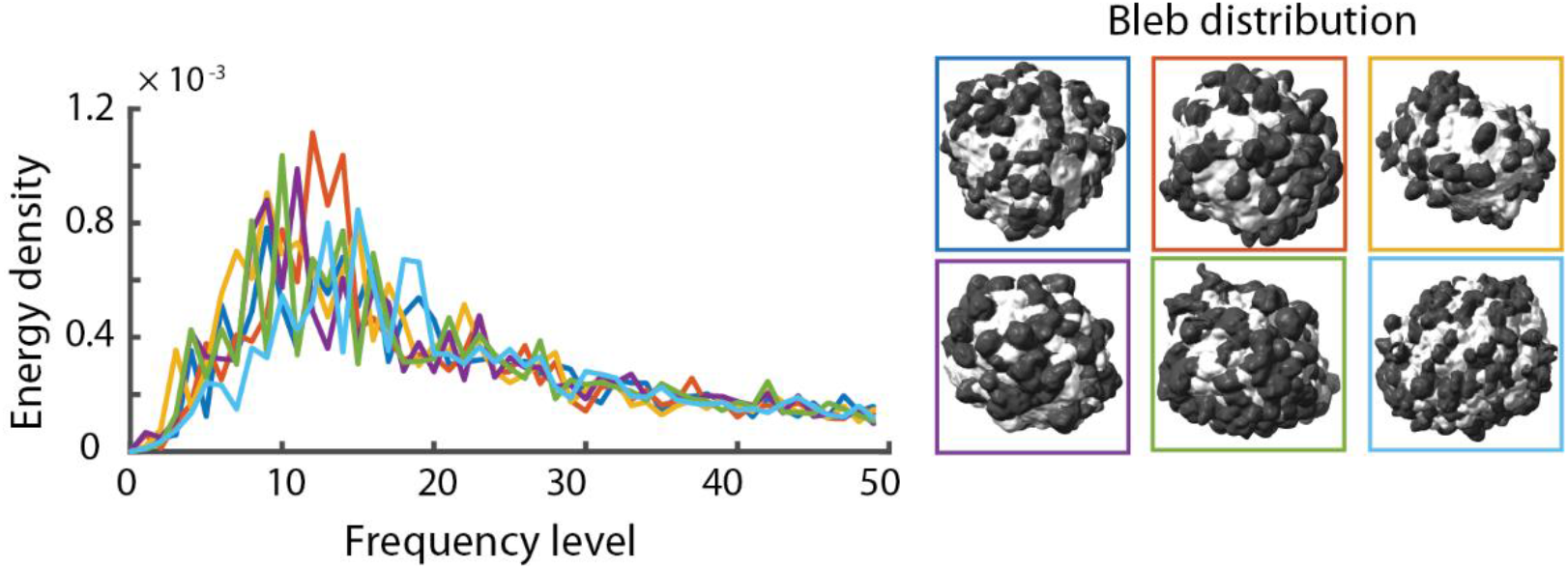
Bleb distribution on melanoma cells. Energy density spectra of the bleb distribution of MV3 melanoma cells expressing NRAS-GFP (left). Surface renderings of the same cells (right) with like colors in correspondence. Blebs are shown dark gray on the cell surface, whereas non-blebs appear white.

**Movie 1. Molecular redistribution over time**. Surface rendering of an MV3 melanoma cell expressing GFP-AktPH, a marker of PI3K activity. The surface is colored by PI3K activity with red indicating regions of high activity and yellow regions of low activity (see the color bar in Fig. 1B).

**Movie 2. LBO eigenvectors at similar frequency levels**. Surface rendering of the LBO eigenvectors *f*_12_, *f*_13_, and *f*_16_ on a sphere (see Fig. S6A).

**Movie 3. LBO eigenvectors with similar numbers of peaks**. Surface rendering of the LBO eigenvectors *f*_12_, *f*_18_, and *f*_24_ on a sphere (see Fig. S6B).

**Movie 4. Molecular organization of tractin-GFP on an MV3 melanoma cell**. Surface rendering of an MV3 melanoma cell expressing tractin-GFP, with surface colored by tractin localization (see Fig 4A).

